# Dorsal Ruffles Enhance Activation of Akt by Growth Factors

**DOI:** 10.1101/324434

**Authors:** Sei Yoshida, Regina Pacitto, Catherine Sesi, Leszek Kotula, Joel A. Swanson

## Abstract

In fibroblasts, platelet-derived growth factor (PDGF) stimulates macropinocytosis and PI 3-kinase (PI3K)-dependent phosphorylation of Akt, leading to activation of mTORC1, a protein complex controlling metabolism and cell growth. PIP_3_, the phosphoinositide product of PI3K that activates Akt, is frequently concentrated within the macropinocytic cups of growth factor-stimulated cells, which suggests that cup structure enhances phosphorylation of Akt by facilitating PI3K activity. However, inhibitors of the cytoskeleton which block cup formation do not reduce Akt phosphorylation in response to high concentrations of PDGF. Because the dynamics of Akt phosphorylation after stimulation by PDGF can differ from those that follow stimulation with epidermal growth factor (EGF), we analyzed the contributions of the actin and microtubule cytoskeleton to activation of Akt by these two growth factors. Actin-rich, circular dorsal ruffles (CDR), analogous to macropinocytic cups, appeared within several minutes of adding EGF or PDGF and often closed to form macropinosomes. Nocodazole, an inhibitor of microtubule polymerization, blocked both PDGF- and EGF-induced CDR formation, and inhibited phosphorylation of Akt in response to EGF but not PDGF. At concentrations that saturate their cognate receptors, EGF stimulated lower maximal levels of Akt phosphorylation than did PDGF. We hypothesized that weak signals elicited by EGF receptors require cytoskeleton-dependent amplification of PI3K for maximal phosphorylation of Akt. In both PDGF- and EGF-stimulated cells, quantitative immunofluorescence showed increased Akt phosphorylation in cells containing CDR, with PIP_3_ and Akt concentrated in CDR and ruffles. Stimulation with low concentrations of PDGF elicited lower levels of Akt phosphorylation, which, like responses to EGF, were inhibited by nocodazole. These results indicate that when receptor signaling generates low levels of PI3K activity, CDR facilitate local amplification of PI3K, PIP_3_ synthesis and phosphorylation of Akt.

## Introduction

Macropinocytosis is a cellular process of solute endocytosis induced by growth factors, chemokines, and various other stimuli (Swanson, 2008, Egami et al., 2014, Buckley and King, 2017, Yoshida et al., 2018). In macrophages, stimulation with macrophage-colony stimulating factor (M-CSF) or the chemokine CXCL12 elicits membrane ruffles, which form cup-shaped structures that close into large endocytic vesicles called macropinosomes (Yoshida et al., 2009, Yoshida et al., 2015a, Pacitto et al., 2017). Macropinosomes either recycle to the plasma membrane or fuse with lysosomes. Stimulation of murine embryonic fibroblasts (MEF) with the growth factors platelet-derived growth factor (PDGF) or epidermal growth factor (EGF) elicits an alternative pathway to cup formation through actin-rich cell surface ruffles, which reorganize to form circular dorsal ruffles (CDR). CDR contract and often close to form macropinosomes (Dubielecka et al., 2010, Schlunck et al., 2004, Lanzetti et al., 2004, Hoon et al., 2012, Itoh and Hasegawa, 2013, Araki et al., 2007, Bryant et al., 2007, Feliciano et al., 2011, Azimifar et al., 2012). CDR and the circular ruffles that comprise macropinocytic cups can localize molecules associated with signal transduction, including phosphatidylinositol 3-kinase (PI3K) and its product phosphatidylinositol (3,4,5)-trisphosphate (PIP_3_) (Yoshida et al., 2009, Yoshida et al., 2015a). Conversely, the formation of CDR or closure of cups into macropinosomes requires PI3K (Wymann and Arcaro, 1994, Araki et al., 1996, Hooshmand-Rad et al., 1997, Valdivia et al., 2017), which suggests that CDR and macropinocytic cups are self-organized structures that require PIP_3_ for complete morphogenesis.

Macropinocytosis provides a mechanism to activate mTORC1 (<u>m</u>echanistic <u>T</u>arget of <u>R</u>apamycin Complex<u>-1</u>), a protein complex that regulates metabolism and cell growth in response to signals generated by growth factors or other ligands at the plasma membrane (Saxton and Sabatini, 2017). mTORC1 is activated at lysosomal membranes by two small GTPases, Rag and Rheb (Saito et al., 2005, Sancak et al., 2010, Betz and Hall, 2013, Saxton and Sabatini, 2017). Macropinosomes induced by receptor activation deliver extracellular nutrients into lysosomes, where lysosome-associated membrane protein complexes detect the increased luminal concentrations of amino acids and trigger the activation of Rag GTPases (Yoshida et al., 2015b, Yoshida et al., 2018, Zoncu et al., 2011). Activated Rag recruits mTORC1 from cytosol to lysosomes (Sancak et al., 2008, Sancak et al., 2010). Additionally, growth factor receptor stimulation of PI3K generates PIP_3_ in plasma membrane, which recruits the serine/threonine kinase Akt via its PH-domain (Manning and Toker, 2017). There, Akt is phosphorylated by PDK1 on Thr308 (pAkt(308)) and by mTORC2 (mTOR complex-2) on Ser473 (pAkt(473) (Ebner et al., 2017a, Ebner et al., 2017b, Zhang et al., 2003). Activated Akt induces the phosphorylation of TSC2, a part of the TSC protein complex which is a GTPase-activating protein (GAP) for Rheb (Potter et al., 2002, Inoki et al., 2002, Inoki et al., 2003, Garami et al., 2003). Phosphorylated TSC complex dissociates from lysosomes, eliminating its GAP activity towards Rheb and thereby permitting Rheb activation of mTORC1 at the lysosomal membrane (Menon et al., 2014).

PIP_3_ generation in CDR or macropinocytic cups may facilitate Akt phosphorylation and downstream signaling. Akt-GFP is recruited to LPS-induced macropinocytic cups in macrophages (Wall et al., 2017) and to macropinosomes induced by active Ras (Porat-Shliom et al., 2008). In response to CXCL12, macrophages expressing fluorescent protein-tagged PH-domain probes show PIP_3_ enriched in membranes of macropinocytic cups (Pacitto et al., 2017), and immunofluorescence staining localizes phosphorylated Akt (pAkt(308)) to macropinocytic cups. These results suggest that the macropinocytic cup provides a signal platform for Akt phosphorylation, upstream of the TSC/Rheb pathway.

Are cups necessary for PI3K-dependent activation of Akt? In macrophages, inhibitors of actin cytoskeleton dynamics or macropinosome formation reduce phosphorylation of Akt in response to CXCL12 (Pacitto et al., 2017). In MDA-MB-231 cells, phosphorylation of Akt in response to the G-protein-coupled receptor (GPCR) lysophosphatidic acid (LPA) requires PI3K catalytic subunit p110β and Rac-dependent macropinocytosis (Erami et al., 2017). In contrast, Akt phosphorylation in response to M-CSF and PDGF, in macrophages and MEF, respectively, is insensitive to cytoskeletal inhibitors (Yoshida et al., 2015b). Single cell analysis of signaling to Akt in response to PDGF and EGF revealed differences in the kinetics and duration of cellular responses (Gross and Rotwein, 2016), which indicated different and heterogeneous cellular responses to the two growth factors and to different concentrations of PDGF. This suggests that qualitative or quantitative features of receptor signaling determine different requirements for cup-restricted PI3K activity.

Here we analyze the contributions of CDR to Akt phosphorylation in response to PDGF and to EGF. We show that CDR enhance the recruitment and phosphorylation of Akt at plasma membrane in MEF, that CDR and macropinosomes formed after stimulation with these growth factors can be inhibited by depolymerization of microtubules, and that CDR facilitate signal amplification when receptor signaling generates lower levels of phosphorylated Akt. The results indicate roles for CDR in the amplification of PI3K and attendant signals.

## Results

### Nocodazole blocks PDGF-induced macropinocytosis and mTORCI activation

To define the time-course of the transition from CDR to macropinosomes in PDGF-stimulated MEF, morphological changes were monitored by time-lapse microscopy (**Fig. 1A**). Following overnight deprivation of MEF for growth factors, CDR were induced within a few minutes of PDGF stimulation (t=1:00, min:sec). CDR then contracted radially, generating macropinosomes (t=6:00). To quantify CDR, cells were fixed at various intervals after PDGF stimulation and the number of the cells with CDR was measured. CDR were induced to maximal levels within a minute of stimulation and decreased afterward (**Fig. 1B**, **top graph**), indicating that induction of CDR by PDGF was an acute response. To study the transition from CDR to macropinosomes, we measured macropinosome formation at the same time intervals (Fig. 1B, **bottom graph**). The number of macropinosome increased over 30 min following stimulation (**Fig. 1B**). Thus, PDGF quickly induce CDR which close into macropinosomes.

**Figure 1.**
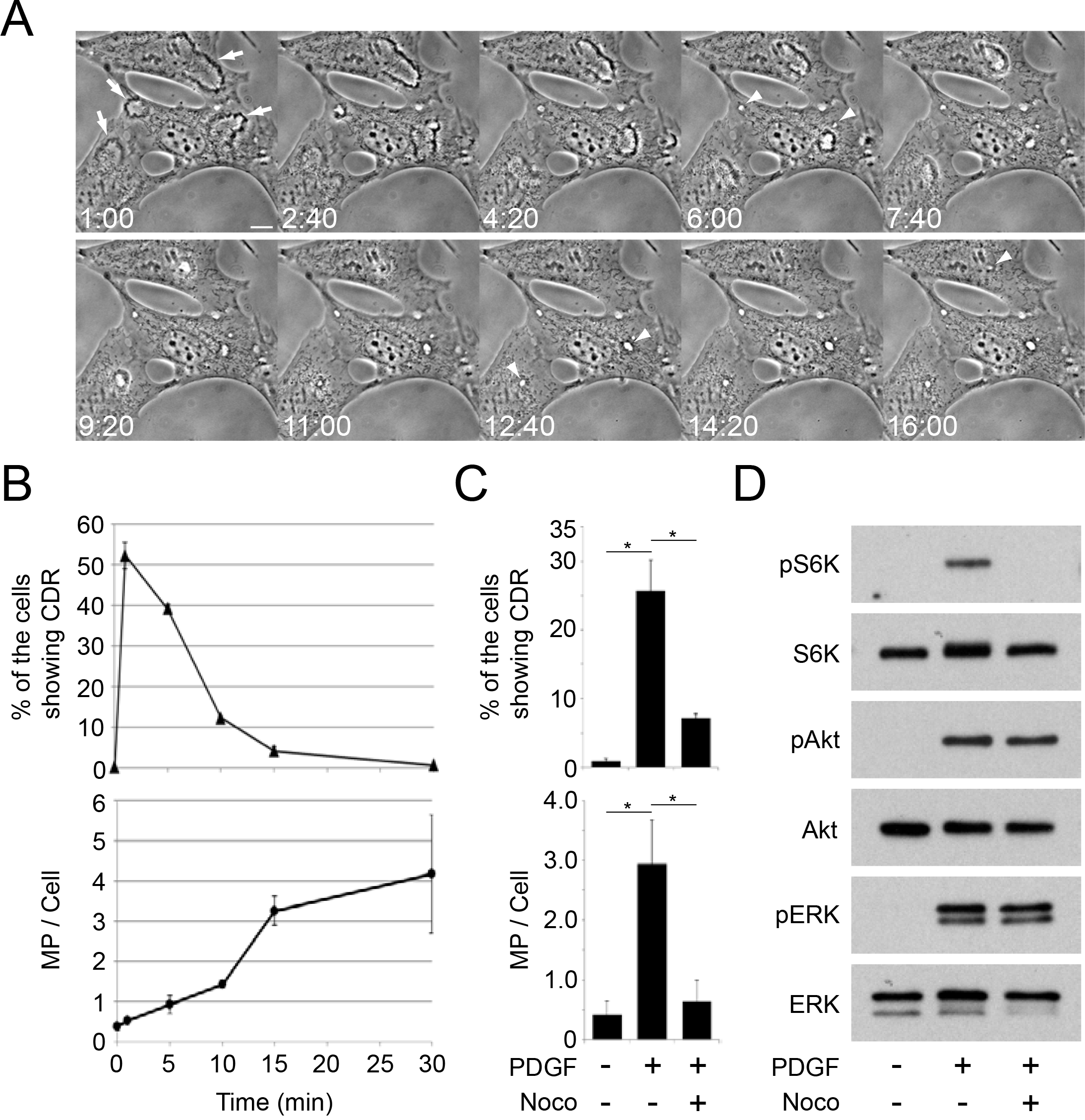
Nocodazole treatment blocks PDGF-induced CDR, macropinocytosis and mTORC1 activation. (**A**) Time-lapse images of MEF stimulated with PDGF (2 nM). CDR were induced within 1 minute (1:00, arrows), which shrank (1:00-4:20), and closed into macropinosomes (arrowheads at 6:00 and 12:40). Scale bar = 10 μm. (**B**) The frequency of cells inducing CDR (top panel) and macropinosomes (bottom panel) at the indicated times after PDGF stimulation. More than 300 cells (top) or 25 cells (bottom) were observed at each time point, and three independent experiments were carried out to calculate the average and standard error. (**C**) Inhibition of CDR and macropinosome formation by nocodazole. Cells were pre-treated with nocodazole (10 μM, 20 min) and stimulated by PDGF for 1 min (for CDR assay) or 15 min (macropinosome assay). (**D**) MEF were incubated in DPBS containing leucine (0.4 mM) for 30 min with/without nocodazole (10 μM, 20 min pre-treatment), then stimulated by PDGF for 15 min. Nocodazole treatment blocked PDGF-induced phosphorylation of S6K but not of Akt or ERK.

Inhibition of cytoskeletal activities by the combination of jasplakinolide and blebbistatin (J/B) blocks PDGF-induced macropinocytosis (Yoshida et al., 2015a). Additionally, we found that the microtubule polymerization inhibitor nocodazole blocked PDGF-induced CDR formation and macropinocytosis (**Fig. 1C**). It was shown previously that inhibition of PDGF-induced macropinocytosis by J/B blocked mTORC1 activation, as measured by S6K phosphorylation (Yoshida et al., 2015b). Likewise, nocodazole blocked PDGF-induced S6K phosphorylation (**Fig. 1D**). Nocodazole treatment did not alter phosphorylation of either Akt or Erk in response to PDGF. These results are consistent with earlier studies showing that PDGF-induced macropinocytosis provides a vesicular pathway for extracellular amino acids to activate mTORC1 (Yoshida et al., 2015b).

### Nocodazole attenuates EGF-induced Akt phosphorylation

To test whether EGF induces CDR and macropinocytosis in MEF, we observed fixed cells by phase contrast microscopy following stimulation for different times (**Figs. 2A** **and B**, **Suppl. Fig. 1A**). Maximal CDR formation was observed after 5 min of stimulation, and decreased during the next 25 min (**Fig. 2A** **and B**). Macropinosome formation increased at 15 min (**Suppl. Fig. 1A**). Nocodazole blocked EGF-induced CDR (**Fig. 2C**) and macropinocytosis (**Suppl. Fig. 1B**). S6K phosphorylation in response to PDGF requires extracellular leucine and is blocked by inhibition of macropinocytosis, indicating that the amount of leucine delivered by macropinosomes to lysosomes determines the magnitude of S6K phosphorylation (Yoshida et al., 2015b). EGF-induced phosphorylation of S6K was similarly dependent on extracellular leucine and macropinocytosis. Although Akt and Erk phosphorylation levels in these conditions were constant after stimulation, the phosphorylation of S6K increased with increasing leucine concentrations (**Fig. 2D**), indicating that EGF-induced mTORC1 activation follows delivery of extracellular leucine into lysosomes. Consistent with a macropinocytosis-mediated delivery of leucine into lysosomes, we observed that nocodazole blocked EGF-induced pS6K (**Fig. 2E**). EGF-induced phosphorylation on threonine 308 of Akt (pAkt(308)) was diminished by nocodazole, while ERK phosphorylation was unaffected (**Fig. 2E**), which suggested that EGF signaling to Akt was enhanced by CDR formation, unlike PDGF. Thus, for both EGF and PDGF, activation of mTORC1 requires macropinocytosis of leucine (vesicular pathway) and activation of Akt (cytosolic pathway). However, inhibition of CDR formation reduced Akt phosphorylation in response to EGF but not PDGF.

**Figure 2.**
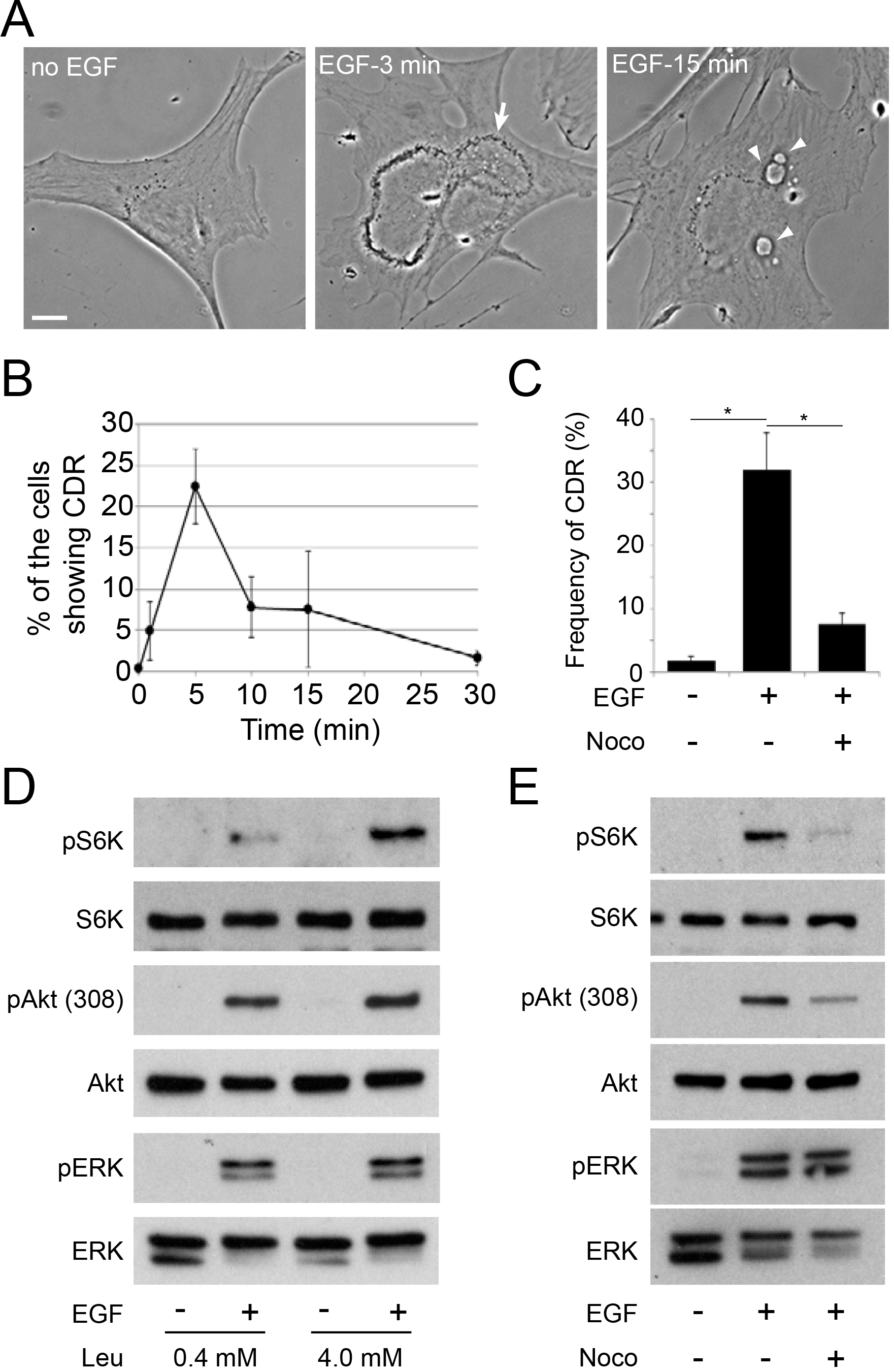
EGF-induced Akt phosphorylation is attenuated by nocodazole treatment. (**A**) CDR (arrow, 3 min) and macropinosome formation (arrow head, 15 min) in response to EGF (16 nM) in MEF. Scale bar = 10 μm. (**B**) The frequency of the cells inducing CDR at the indicated times after EGF stimulation. (**C**) Nocodazole treatment blocked EGF-induced CDR formation. (**D**) Effects of leucine concentration on EGF-induced S6K phosphorylation. Cells were incubated in DPBS containing 0.4 mM or 4.0 mM leucine for 30 min, then stimulated by EGF for 15 min. (**E**) Effects of nocodazole treatment on EGF-induced signal pathways. Cells were incubated in DPBS containing 0.4 mM leucine for 30 min with or without nocodazole, then stimulated by EGF for 15 min. Nocodazole treatment inhibited S6K phosphorylation and attenuated Akt phosphorylation without affecting ERK phosphorylation.

### Inhibition of CDR formation attenuates low-level phosphorylation of Akt

We reported earlier that pAKT(308) induced by CXCL12 in macrophages was blocked by inhibition of macropinocytic cup formation, and that the magnitude of pAKT(308) generated in response to CXCL12 was less than that elicited by M-CSF (Pacitto et al., 2017). Based on that study, we hypothesized that the different inhibitory effects of nocodazole on levels of pAkt(308) in response to PDGF and EGF were related to the different magnitudes of Akt phosphorylation elicited by the two growth factors. To test this, we measured the phosphorylation of Akt (pAkt(308) and pAkt(473)) induced by different doses of EGF or PDGF. When cells were stimulated with concentrations of EGF and PDGF that saturate receptors (Ozcan et al., 2006, Fretto et al., 1993), the level of Akt phosphorylation induced by EGF was less than that induced by PDGF. Notably, the magnitude of ERK phosphorylation (pERK) under each experimental condition was the same (**Fig. 3A**). Similar to the effects of nocodazole, inhibition of the actin cytoskeleton by jasplakinolide and blebbistatin (J/B) decreased Akt phosphorylation in response to EGF, without affecting ERK phosphorylation (**Fig. 3B**). Akt phosphorylation induced by higher concentrations of EGF (160 nM) was also attenuated by nocodazole treatment (**Fig. 3C**). Thus, EGF induced lower maximal levels of Akt phosphorylation than did PDGF, and those low levels of phosphorylation were sensitive to cytoskeletal inhibition.

**Figure 3.**
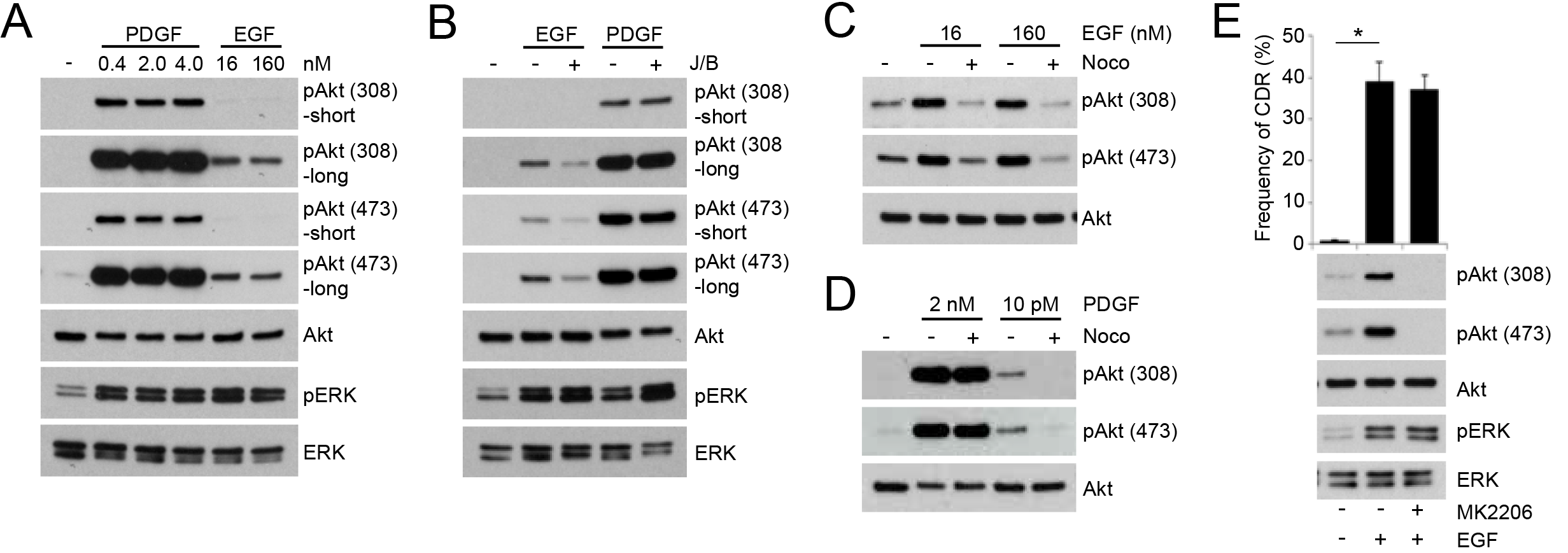
Inhibition of CDR attenuates Akt phosphorylation in response to stimuli which generate weaker signals. (**A**) Comparison of the magnitudes of Akt phosphorylation induced after 15 min stimulation by PDGF (2 nM) and EGF (16 nM). Intensities of pAkt(308) and pAkt(473) were stronger in response to PDGF than to EGF. (**B**) Jasplakinolide and blebbistatin (J/B) attenuated pAkt(308) and pAkt(473) responses induced by EGF (16 nM), but not by PDGF (2 nM). (**C**) Nocodazole treatment attenuated pAkt(308) and pAkt(473) induced by EGF. (D) Nocodazole treatment attenuated pAkt(308) and pAkt(473) induced by 10 pM PDGF. (**E**) The Akt inhibitor MK2206 did not block EGF-induced CDR formation.

The differential sensitivities of Akt phosphorylation to cytoskeletal inhibition in response to PDGF and EGF could be due to qualitative differences in signaling between the two growth factor receptors, which render EGF receptor signaling to Akt sensitive to cytoskeletal inhibition. Alternatively, the sensitivity to these inhibitors could reflect a shared mechanism of Akt signal amplification which is only apparent when signals are low. To address this possibility, we used low concentrations of PDGF (10 pM) to elicit lower levels of Akt phosphorylation, and observed that nocodozole treatment attenuated pAkt(308) and pAkt(473) responses (**Fig. 3D**). The Akt-specific inhibitor MK2206 blocked synthesis of both pAkt(Thr308) and pAkt(Ser473), but did not affect CDR formation or ERK phosphorylation (**Fig. 3E**), indicating that Akt functions downstream of CDR. Together, these results indicate that cytoskeletal activities facilitate Akt phosphorylation in response to weaker stimuli.

### CDR correlate with increased Akt phosphorylation

We developed quantitative immunofluorescence methods to determine the magnitudes of Akt phosphorylation in individual cells (Pacitto et al., 2017). To determine the applicability of this method for MEF, cells were stimulated with 2 nM PDGF for 15 minutes, with or without inhibitors for PI3K catalytic subunits p110α (A66) (Jamieson et al., 2011) or p110β (TGX-221) (Jackson et al., 2005), then samples were fixed and pAkt(308) and Akt were labeled for immunofluorescence microscopy (**Suppl. Fig. 2A**). The ratio images (pAkt/Akt) were calculated and the intensities were quantified relative to unstimulated cells. The ratio values of the PDGF-stimulated samples were higher than those of controls, and were reduced by the inhibitor treatments (**Suppl. Figs. 2A** **and B**). Cell lysates were prepared under the same experimental conditions and pAkt(308) was measured by western blot analysis. The results indicate that quantitative ratiometric immunofluorescence of pAkt and Akt in individual cells reflected the results of western blot analysis of cell lysates (**Suppl. Fig. 2C**).

Using this method, we investigated the relationship between CDR and Akt phosphorylation in single cells. After stimulation with growth factors, cells were labelled for immunofluorescence microscopy, and the relative intensities of pAkt(308) to Akt were measured ratiometrically. Based on the combinations of ligands and whether cells formed CDR, five different conditions were measured: no growth factor treatment, EGF/no CDR, EGF/with CDR, PDGF/no CDR, and PDGF/with CDR. Analysis of phase contrast and ratio images showed that, after EGF treatment, ratio values were higher in cells with CDR than in cells lacking CDR (**Figs. 4A** **and B**). The ratio values of EGF/no CDR were significantly higher than those with no growth factor treatment, indicating that CDR formation was not required for pAkt(308) (**Figs. 4A** **and B**). We also found that the ratio values of PDGF-stimulated cells with CDR were slightly higher than in PDGF-stimulated cells lacking CDR (**Fig. 4A**); however, the differences were not statistically significant (**Fig. 4B**). Similar relationships were observed for pAkt(473) (**Suppl. Fig. 3**).

**Figure 4.**
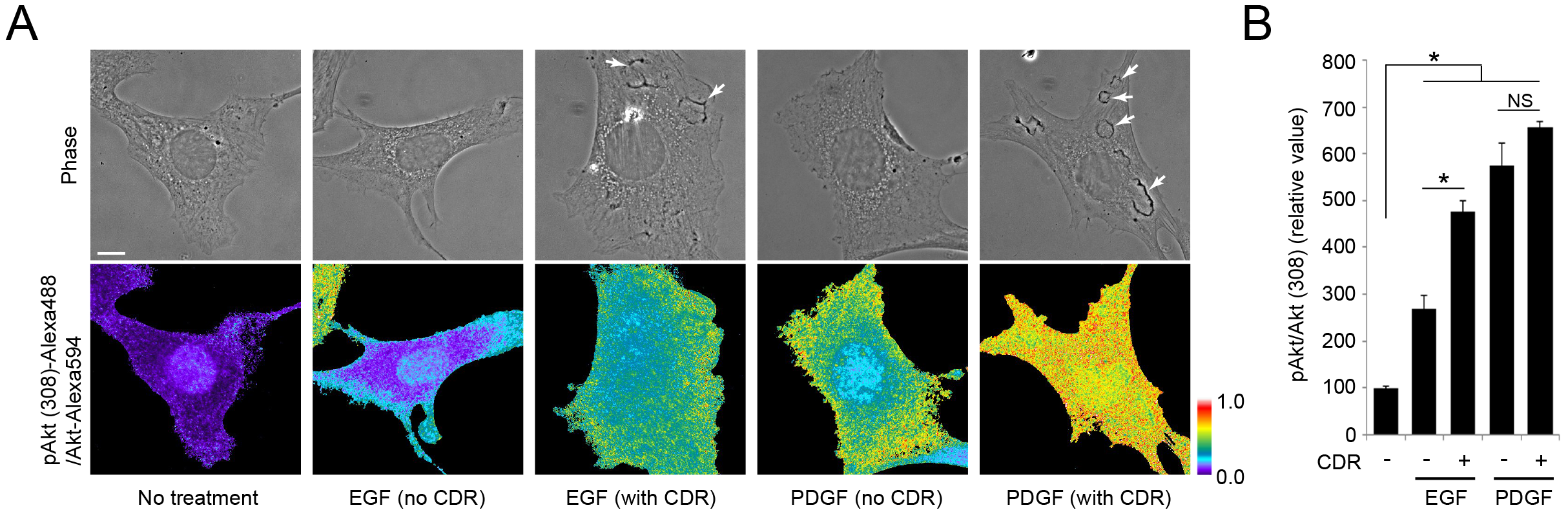
Correlation between the formation of CDR and the magnitude of Akt phosphorylation. (**A**) Phase and ratio (pAkt (308)/Akt) images for MEF in different conditions. Cells were treated with or without EGF (16 nM) or PDGF (2nM) for 3 min. After fixation, samples were stained with anti-pAkt(308) and anti-Akt antibodies. Phase contrast, pAkt(308) and Akt images were taken, then the ratio images were obtained by dividing the pAkt(308) image by the Akt image. Arrows indicate CDR. Scale bar = 10 μ.m. Color bar indicates relative values of ratio intensities. (**B**) Quantification of ratiometric imaging in (A). The relative ratio value of EGF-stimulated cells with CDR (EGF, CDR+) was significantly higher than that of EGF-stimulated cells without CDR (EGF, CDR-). More than 10 cells were observed for each assay. A one-way ANOVA was applied for the statistics. *p<0.05.

As observed by western blot analysis (**Fig. 3C** **and D**), the quantitative immunofluorescence analysis also determined that nocodazole attenuated pAkt(308)/Akt ratios in EGF-stimulated cells (**Fig. 5A**). Likewise, the ratios in 10 pM PDGF-stimulated cells were lower than those from 2 nM PDGF, and were reduced by nocodazole treatment (**Fig. 5B**), confirming that Akt phosphorylation in response to low levels of stimulation was sensitive to nocodazole.

**Figure 5.**
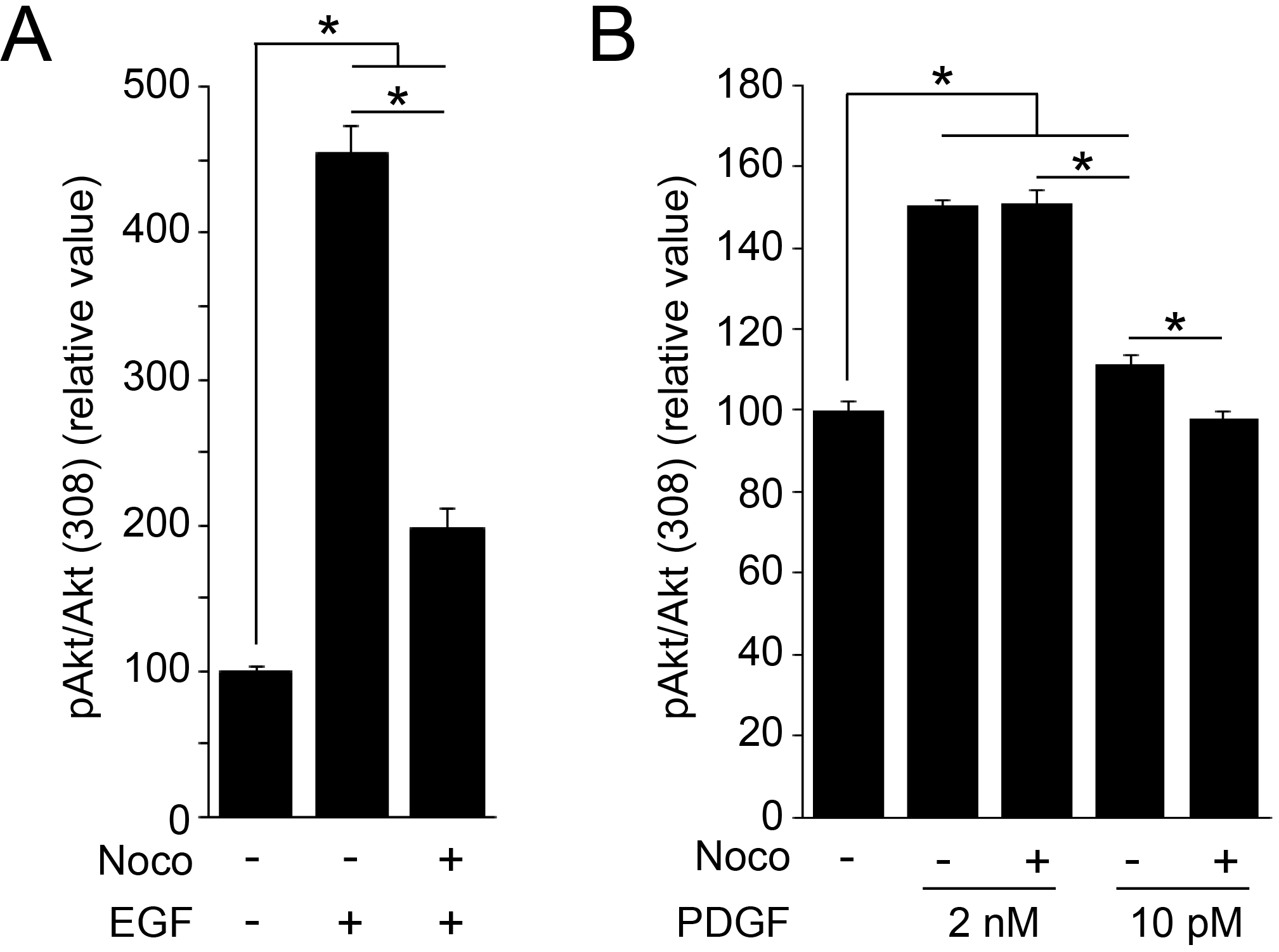
Nocodazole reduces phosphorylation of Akt. (**A**) MEF were stimulated by EGF (16 nM) for 3 min with or without nocodazole pre-treatment. After fixation, samples were stained with anti-pAkt(308) and anti-Akt antibodies, and phase, pAkt(308), and Akt images were taken. Ratio images were calculated to quantify the intensity of pAkt in each condition. The relative ratio values of EGF-stimulated cells were significantly higher than those without treatment and were reduced by nocodazole pre-treatment. More than 15 cells were observed for each condition. A one-way ANOVA was applied for the statistics. *p<0.05. (**B**) MEF were stimulated by PDGF (2 nM or 10 pM) for 3 min with or without nocodazole pre-treatment. Ratio images of pAkt (308) and Akt were calculated to quantify the intensity of pAkt in each condition. The relative ratio value of 10 pM PDGF-stimulated cells was significantly higher than that of no treatment and was attenuated by nocodazole. The values for cells stimulated with 2 nM PDGF were not attenuated by nocodazole. More than 15 cells were observed for each condition. A one-way ANOVA was applied for the statistics. *p<0.05.

### PIP_3_, Akt and phospho-Akt are enriched in CDR and cell ruffles

Akt activation requires recruitment to the plasma membrane via the interaction of the PH domain of Akt with PIP_3_ or PI(3,4)P_2_ (Ebner et al., 2017a). To observe distributions of PIP_3_ during CDR formation, we performed ratiometric imaging of yellow fluorescent protein (YFP)-tagged Btk-PH, a probe for PIP_3_ (Varnai et al., 1999) and free cyan fluorescence protein (CFP). After stimulation with PDGF or EGF, YFP-Btk-PH recruitment to ruffles and CDR was observed in living and fixed cells (**Fig. 6**). The results suggested that Akt should preferentially bind plasma membrane inside CDR. To localize Akt, MEF expressing free CFP were stimulated 3 min with EGF or PDGF, then were fixed and labeled for immunolocalization of Akt. Ratiometric imaging of Akt-Alexa594 and CFP, which corrects for variations in cell thickness (Swanson, 2002), indicated higher concentrations of Akt within CDR (**Fig. 7 A**), which was confirmed by quantitative analysis (**Fig. 7 B**). Similarly, PDGF induced CDR in which the relative value of Akt-Alexa594/CFP was higher than the average for the entire cell (**Fig. 7 A** **and B**). These results indicate that Akt was recruited to CDR by the local generation of PIP_3_ inside the structure. Next, we observed the distributions of pAkt(308) and pAkt(473) in EGF-stimulated MEF. Cells expressing CFP were stimulated by EGF for 3 minutes, then fixed and prepared for ratiometric immunofluorescence microscopy of pAkt(308) or pAkt(473). The ratio values of pAKT(308)/CFP and pAkt(473)/CFP were higher inside CDR than the cell average, although high values were also observed in ruffle-rich regions outside CDR (**Fig. 7 C** **and D**). Together, the quantitative imaging supports a model in which CDR create domains of plasma membrane that support PI3K-dependent phosphorylation of Akt.

**Figure 6.**
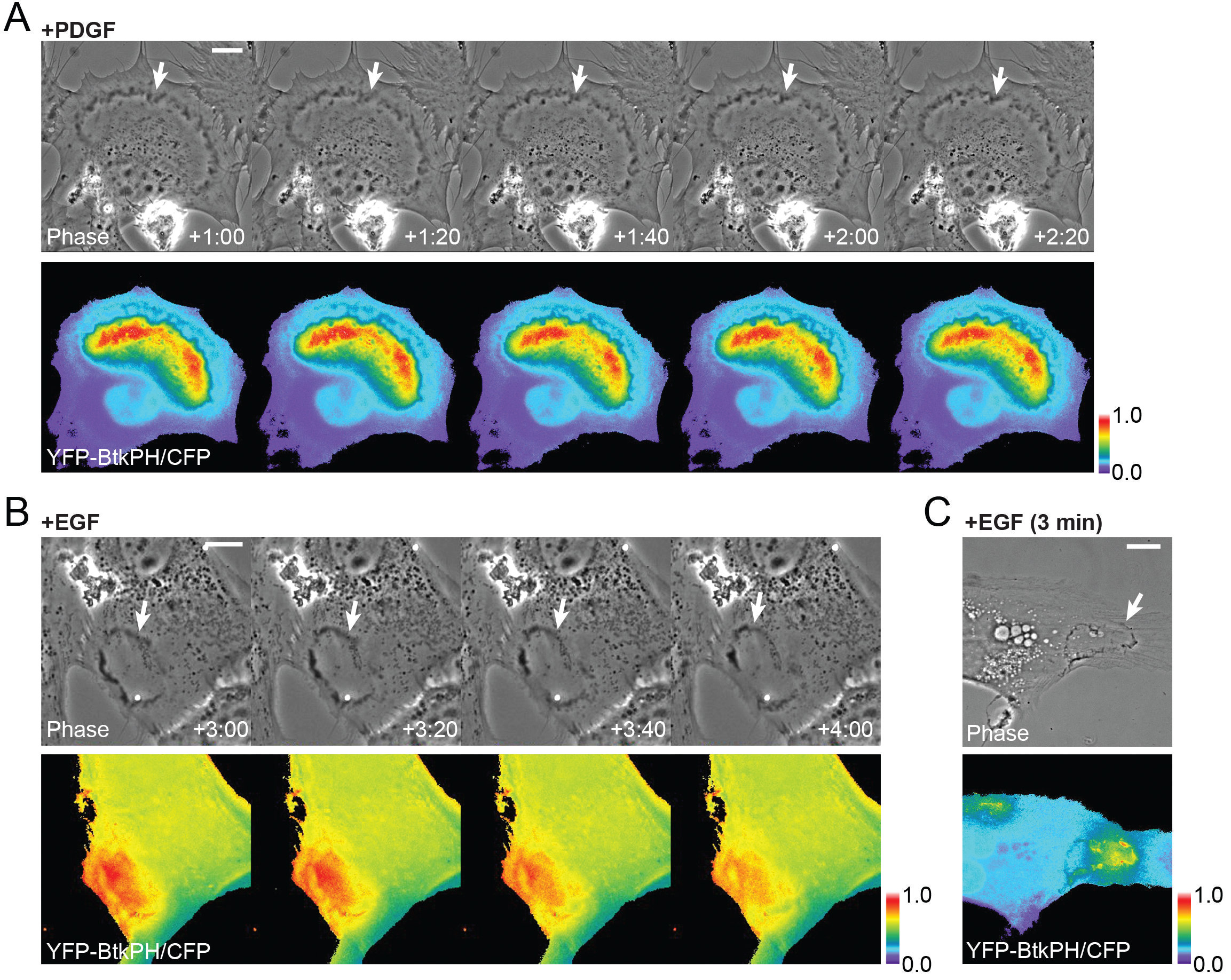
Btk-PH recruitment inside CDR. (**A, B**) Live-cell imaging of MEF expressing YFP-Btk-PH, a probe protein for PIP_3_, and CFP as a reference after stimulation by PDGF (A) or EGF (B). Comparison of phase contrast and ratio (YFP-Btk-PH/CFP) images shows high ratio values inside CDR (arrow). Time after addition of growth factor is indicated at bottom of phase contrast images (minutes : seconds). Scale bar = 10 μm. Color bar indicates relative value of ratio intensities. (**C**) MEF expressing YFP-Btk-PH and CFP were fixed after 3 min stimulation with EGF. High ratio values were observed inside CDR (arrow). Scale bar = 10 μm. Color bar indicates relative value of ratio intensities.

**Figure 7.**
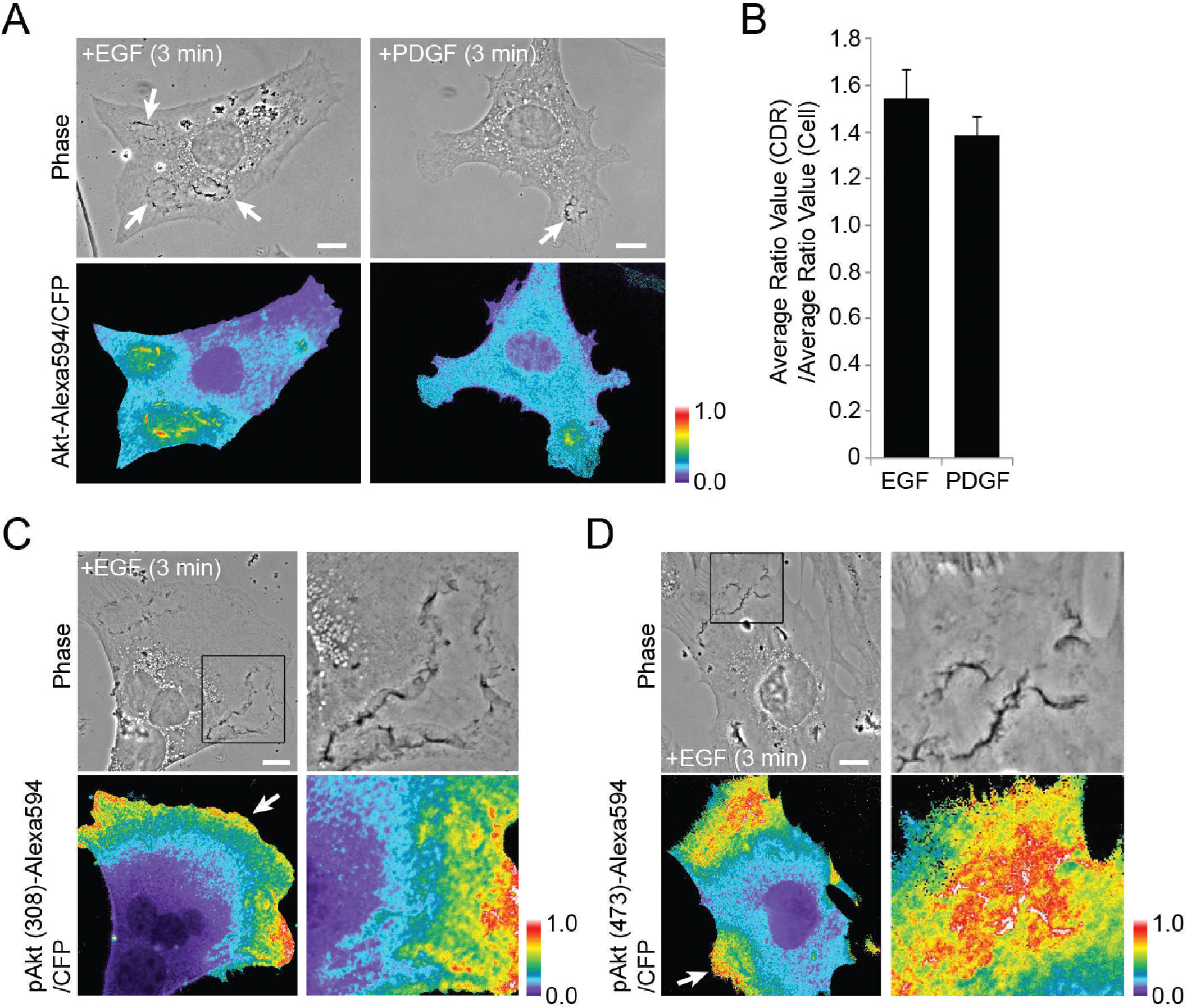
Characterization of CDR as platforms for Akt signaling. (**A**) Fluorescence microscopy of Akt recruitment to CDR in MEF. Cells expressing CFP were stimulated with EGF or PDGF for 3 min and fixed for immunofluorescence staining for Akt (Alexa594-tagged secondary antibody). Comparisons of phase contrast and ratio images (Akt-Alexa594/CFP) show higher ratio values in CDR (arrows). (**B**) Quantification of ratio images of (A). The ratio value inside CDR is divided by that of whole cell, and the average and SE were calculated (13 or 9 images of EGF or PDGF-stimulated MEF were observed, respectively). (**C, D**) Fluorescence microscopy observations were used to observe EGF-induced pAkt(308) (C) or pAkt (473) (D). Strong signals were observed both inside (cropped images) and outside (arrow) of CDR. Scale bar = 10 μm. Color bar indicates relative value of ratio intensities.

## Discussion

This study identifies a role for CDR in amplifying growth factor receptor signaling to Akt. Live-cell imaging and immunofluorescence localized PIP_3_ inside CDR in MEF. Western analysis showed that inhibition of cytoskeletal activities limited Akt phosphorylation in response to EGF or to low concentrations of PDGF, but not to higher concentrations of PDGF (**Fig 3. C** **and D**). Quantitative analysis of immunofluorescence staining demonstrated greater levels of Akt phosphorylation in cells with CDR than in cells without (**Fig. 4**, **Suppl. Fig. 3**). Thus, as restricted areas of PIP_3_ accumulation, CDR provide platforms for Akt phosphorylation. We propose that CDR amplify Akt phosphorylation when receptor signaling generates low levels of PI3K activity, either by low concentrations of ligand or by innate differences in receptor signaling to PI3K (**Fig. 8**).

**Figure 8.**
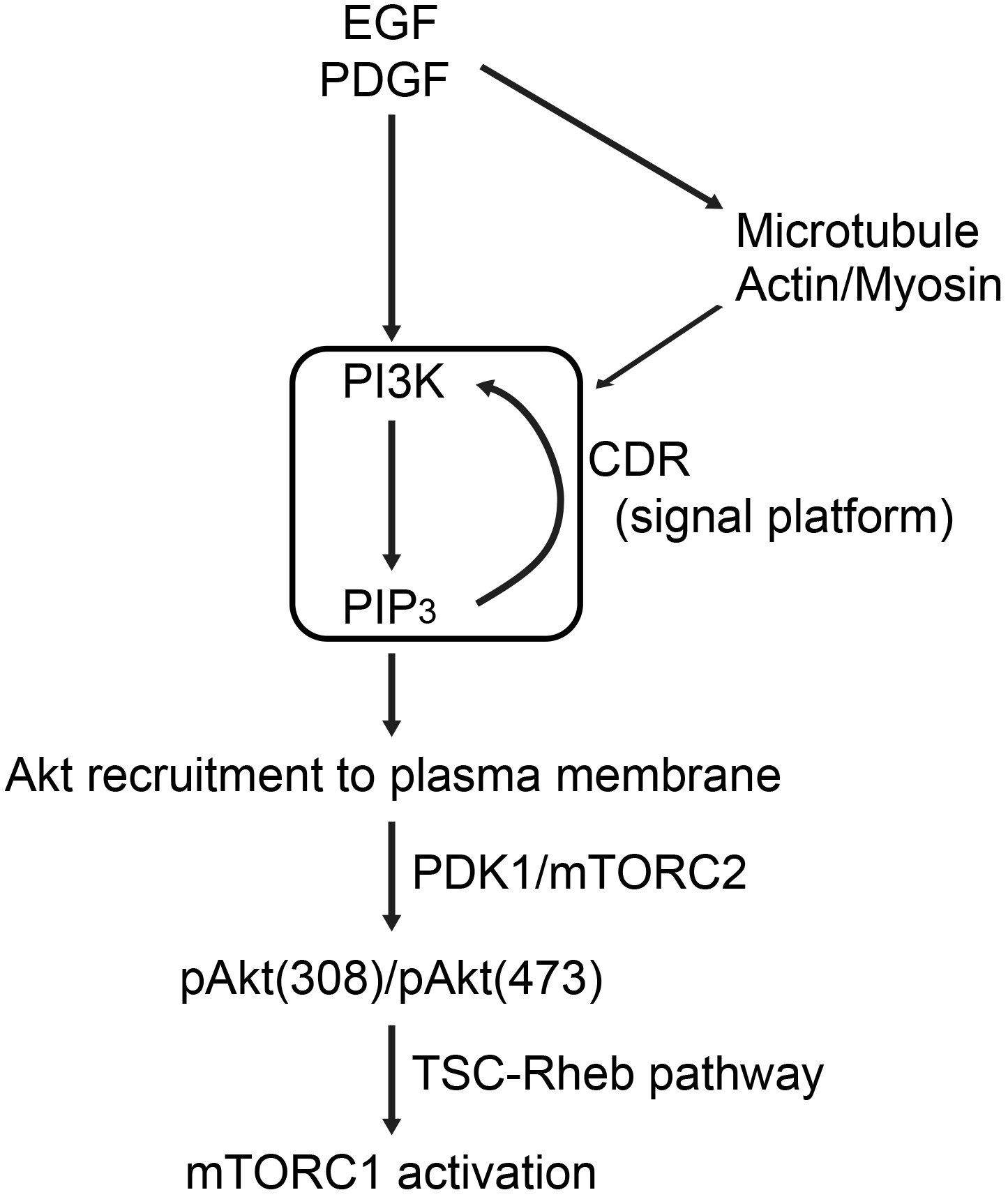
A model for CDR-dependent Akt phosphorylation. CDR induced by EGF or PDGF provide platforms for Akt signal amplification. After stimulation, CDR are formed by cytoskeletal activities. Meanwhile, activated PI3K generates PIP_3_ at the plasma membrane. The molecular mechanism of PIP_3_ restriction to CDR is still unknown, but could be due to a positive feedback between PIP_3_ and PI3K, supported by diffusion barriers in CDR. PIP_3_ concentrations inside CDR recruit Akt to the plasma membrane. Akt is phosphorylated on Thr308 and Ser473 by PDK1 and mTORC2, respectively, upstream of the TSC-Rheb pathway for mTORC1 activation.

How does Akt signal amplification localize to CDR or macropinocytic cups? Structural barriers in cup margins limit lateral diffusion of membrane-tethered proteins within the inner leaflet of cup membranes (Welliver et al., 2011). Such barriers should confine phospholipids or the enzymes that modify them within CDR and cups, consequently localizing signal cascades to these regions of plasma membrane (Welliver and Swanson, 2012). PIP_3_ synthesized by PI3K within CDR may be retained there by the diffusion barrier, allowing positive feedback stimulation of PI3K by PIP_3_, which could be mediated by Ras, Rac or the adapter protein Gab1 (Rodrigues et al., 2000, Castellano and Downward, 2011, Fritsch et al., 2013, Ding et al., 2015). The elevated concentrations PIP_3_ within CDR could also enhance barrier formation through activation of Rac1 or Ras (Kay et al., 2018, Delos Santos et al., 2015, Hoon et al., 2012, Lien et al., 2017). If PIP_3_ in membranes must exceed some concentration threshold to allow binding of Akt, PDK1 or mTORC2, or if ruffles or cups somehow enhance coupling between PDK1 and Akt, then activation of PI3K in CDR could provide a mechanism to amplify Akt phosphorylation.

Unlike EGF and low concentrations of PDGF (10 pM), higher concentrations of PDGF (2-4 nM) induced Akt phosphorylation that was independent of CDR. Western blotting (**Fig. 3**) and microscopic assays (**Fig. 4**) showed that at receptor-saturating concentrations, PDGF induced stronger Akt phosphorylation than did EGF. Although EGF elicited lower maximal levels of Akt phosphorylation than did PDGF, levels of ERK phosphorylation by the two growth factors were comparable. This suggests that receptors for EGF and PDGF initiate different overall levels of PI3K activity. A similar relationship appears in macrophages, where CXCL12 generates low levels of pAkt(308) which can be reduced by inhibition of macropinocytic cup formation, and M-CSF generates higher levels of pAkt(308) which are not reduced by such inhibition. These results indicate that CDR in MEF and macropinocytic cups in macrophages provide a mechanism to boost Akt phosphorylation when PI3K generates relatively less PIP_3_ (**Fig. 8**).

The kinetics of Akt phosphorylation differ in response to different ligands and to different concentrations of ligands (Gross and Rotwein, 2016). The timing of CDR formation corresponds to the transient activation of Akt by EGF and PDGF and may reflect a transient amplification of PI3K within CDR. Sustained activation of Akt observed in response to high concentrations of PDGF (Gross and Rotwein, 2016) may reflect a cytoskeleton-independent mechanism of PIP_3_ generation.

This and previous studies of signaling to Akt (Yoshida et al., 2015b, Pacitto et al., 2017, Erami et al., 2017) support a concept in which cells amplify weak PI3K signals using confined subregions of plasma membrane that limit lateral diffusion of PIP_3_ in the membrane (Kay et al., 2018). Cytoskeleton-dependent amplification of Akt signaling may reflect the normal behavior of growing cells, in which constant concentrations of growth factors trigger stochastic, localized signal cascades (Yoshida et al., 2009, Yoshida et al., 2018). Cytoskeleton-independent signaling could be an aberrant consequence of receptor overexpression or dysregulation of PI3K.

## Materials and Methods

### Reagents, plasmids and antibodies

Recombinant murine PDGF-BB and recombinant murine EGF were from Peprotech. Fluorescein isothiocyanate-labeled dextran, avg. mol. wt. 70,000 (FDx70) was from Molecular Probes. Nocodazole and Blebbistatin were from Abcam. Jasplakinolide was from Enzo Life Science. MK 2206 was from ApexBio. A66 and TGX 221 were from Symansis. Anti-phosho-S6K(Thr389) (#9234), anti-S6K (#2708), anti-phospho-Akt(Thr308) (#4056), anti-phospho-Akt(Ser473) (#4060), anti-Akt (#9272), anti-phospho-ERK1/2(Thr202/Tyr204) (#4376), and anti-ERK1/2 (#4695) antibodies for western blot analysis were from Cell Signaling. Anti-phospho-Akt(Thr308) (#2695), anti-phospho-Akt(Ser473) (#4060), and anti-Akt (#2920) for immunofluorescence staining were from Cell Signaling. HRP-conjugated goat anti-rabbit IgG was from GE Healthcare. Anti-rabbit Alexa 488 and anti-mouse Alexa 594 antibodies were from Invitrogen. Plasmids pECFP-N1 for free CFP and pmCitrine-BtkPH-N1 for YFP-Btk-PH were described previously (Yoshida et al., 2009). FuGENE HD was from Promega.

### Cell culture, inhibitor treatment, and transfection

Mouse embryonic fibroblasts (MEF) were cultured as described previously (Yoshida et al., 2015b, Gupta et al., 2007). Briefly, cells were cultured in Dulbecco’s Modified Eagle Medium (DMEM, Life Technologies 11995) with 10% fetal calf serum (FCS) and penicillin/streptomycin. For inhibitor treatments, cells were pretreated with nocodazole (10 μM, 20 min), MK2206 (2 μM, 30 min), A66 (3 μM, 30 min), and TGX 221 (0.5 μM, 30 min) in low-glucose DMEM (Life Technologies 11885). A combination of blebbistatin (75 μM for 35 min) and jasplakinolide (1 μM for 15 min) was also used. FuGENE HD was used for transfection according to the manufacturer’s protocol.

### Microscopy and live-cell imaging

A Nikon Eclipse TE-300 inverted microscope with a 60x numerical aperture 1.4, oil-immersion PlanApo objective lens (Nikon, Tokyo, Japan) and a Lambda LS xenon arc lamp for epifluorescence illumination (Sutter Instruments, Novato, CA) were used to collect phase-contrast and fluorescence images. Fluorescence excitation and emission wavelengths were selected using a 69008 set (Chroma Technology, Rockingham, VT) and a Lambda 10-2 filter wheel controller (Shutter Instruments) equipped with a shutter for epifluorescence illumination control. A Photometrics CoolSnap HQ cooled CCD camera (Roper Scientific, Tucson, AZ) was used for recording. For live-cell imaging, cells plated onto glass-bottom, 35 mm diameter dishes (MatTek Corp.) were cultured in low-glucose DMEM overnight. Cells were stimulated with 2 nM PDGF or 16 nM EGF, and imaged at 20-sec intervals. Time-lapse images were processed using MetaMorph software.

### Circular Dorsal Ruffle assay

MEF were cultured on coverslips in low-Glucose DMEM without FBS overnight. Cells were stimulated with 2 nM PDGF or 16 nM EGF (1-3 minutes), then fixed with fixation buffer 1 (20 mM HEPES, ph 7.4, 4% paraformaldehyde, 4.5% sucrose, 70 mM NaCl, 10 mM KCl, 10 mM MgCl2, 2 mM EGTA, 70 mM lysine-HCl, 10 mM sodium periodate) at room temperature for 15 minutes. The fixed cells were washed with washing buffer (Tris buffered saline (TBS): 20 mM Tris-HCl, pH 7.4, 150 mm NaCl, 4.5% sucrose) for three times 10 minutes at room temperature and mounted for microscopy. The frequency of the cells inducing CDR was determined from images of more than 300 cells per condition. Cells were randomly selected and the number of the cells with CDR was counted. The frequency was calculated as (number of the cells with CDR)/(total number of cells observed). The average and standard error of the frequencies were calculated from three independent experiments. One-way ANOVA was applied for the statistics.

### Macropinosome assay

Macropinosome assays were performed as described previously (Yoshida et al., 2015b). Cells were cultured on coverslips in low-glucose DMEM without FBS overnight. PDGF (2 nM)/EGF (16 nM) and FDx70K (1 mg/ml) were added and the cells were incubated 15 minutes at 37 C. Cells were fixed with fixation buffer2 (20 mM HEPES, pH 7.4, 2% paraformaldehyde, 4.5% sucrose, 70 mM NaCl, 10 mM KCl, 10 mM MgCl2, 2 mM EGTA, 70 mM lysine-HCl, 10 mM sodium periodate) at 37 C for 30 minutes following three washes with warmed DPBS to remove extracellular FDx70. The fixed cells were washed with washing buffer for three times 10 minutes at room temperature and mounted. Phase-contrast and FDx70 images of each sample were taken and merged after subtracting background signal using MetaMorph v6.3 (Molecular Devices). More than 25 cells were observed, and the number of induced macropinosomes per cell was determined by counting FDx70-positive vesicles on the merged images. If the signal of the FDx70-positive structure on the merged images was not clearly a macropinosome, then both FDx70 and phase images were observed separately to confirm macropinosome identity. The average number of induced macropinosomes per cell was calculated based on the results from three independent experiments. One-way ANOVA was applied for the statistics.

### Immunofluorescence staining of Akt and pAkt

Immunofluorescence staining of Akt and pAkt was carried out as described previously (Pacitto et al., 2017). MEF, treated for 3 or 15 min, with or without PDGF or EGF, were fixed at room temperature for 10 min with fixation buffer 1. Cells were washed with TBST buffer (50 mM Tris, 150 mM NaCl, 0.1 % Tween 20, pH 7.6) and permeabilized in freshly prepared 0.2% saponin in TBS (w/v) for 15 min at room temperature, and then incubated in 1% BSA in TBST as the blocking buffer, for 30 min at room temperature. Anti-Akt and anti-pAkt (Thr308)/anti-pAkt (Ser 473) antibodies were diluted at 1:50 in blocking buffer and incubated with the samples for 2 h at room temperature as primary antibody treatment. Samples were washed with TBST three times 10 minutes. Anti-rabbit Alexa 488 and anti-mouse Alexa 594 antibodies were diluted at 1:200 in blocking buffer and incubated with the samples for 1.5 h at room temperature as secondary antibody treatment. Samples were washed with TBST for 10 minutes three times, and mounted.

### Quantification of pAkt/Akt ratio image

Quantification of pAkt/Akt ratio image was carried out as described previously (Pacitto et al., 2017). pAkt and Akt images (excitation filter: 500/20-emission filter: 535/30, excitation filter: 572/35-emission filter: 535/30, respectively) were corrected for shade, bias, and background (Pacitto et al., 2017, Hoppe, 2012). A binary image map was then created from the corrected Akt (denominator) image. The corrected Akt images and binary images were combined using the “Logical AND” command of MetaMorph, and the resulting image was saved as “AND image”. The corrected pAkt image was divided by the AND image, multiplied by 100, and saved as “Ratio image”. These Ratio images were used to quantify the average pAkt/Akt ratio value in single cells. Ratio images were thresholded to exclude background areas and cellular regions were selected manually. With the “Region Measurements” command of MetaMorph, the Integrated Intensity of pAkt/Akt ratio and Threshold Area for a target cell were logged to Excel.

In Excel, integrated intensities were divided by the threshold areas to yield relative intensities as the average pAkt/Akt ratio value in a single cell. More than 10 different cells (**Fig. 4** **and** **Suppl. Figs. 2** **and 3**) or 20 different cells (**Fig. 5**) were observed in each condition. A one-way ANOVA was applied for the statistics.

### Ratio imaging and quantitative analysis of fixed CFP-expressing cells

MEF expressing CFP were treated with EGF or PDGF for 3 min. Cells were fixed and stained with Akt or pAkt antibody with Alexa-594-tagged secondary antibody. Phase, CFP, Akt or pAkt images were taken. Alexa-594/CFP ratio images were generated after shade-bias-background correction. Cell area and CDR area(s) were determined on the phase image using the drawing tool of MetaMorph, and the confined regions were transferred to the ratio image by MetaMorph command. The average ratio value inside the Cell areas and CDR areas were measured by applying the Region Measurement tool. The results were logged to the Excel sheet and (Average Ratio Value at CDR) / (Average Ratio Value of whole Cell) were calculated. For the quantification (**Fig. 7B**), 13 or 9 cell images were observed of EGF-or PDGF-stimulated MEFs, respectively. Student’s t-test was applied for the statistics.

### Ratiometric live-cell imaging

Plasmids pECFP-N1 and pmCitrine-BtkPH-N1 were used for free CFP and YFP-Btk-PH, respectively. Plasmids were transfected into MEFs by Fugene HD according to the manufacturer’s protocol. After PDGF/EGF stimulation, phase-contrast, YFP and CFP images of live MEFs were captured every 20 s for 10 min. Ratio images of YFP-Btk-PH relative to CFP were generated, as described previously (Yoshida et al., 2015b, Pacitto et al., 2017).

### Cell lysates and western blotting

Cells were pretreated with the inhibitors and stimulated with PDGF for 15 minutes, then were lysed for 10 min in cold lysis buffer (40 mM HEPES, pH 7.5, 120 mM NaCl, 1 mM EDTA, 10 mM pyrophosphate, 10 mM glycerophosphate, 50 mM NaF, 1.5 mM Na3Vo4, 0.3% CHAPS, and a mixture of protease inhibitors (Roche Applied Science)) (Yoshida et al., 2011). Lysates were centrifuged at 13,000xg for 15 min at 4 C. The supernatant was mixed with 4x SDS sample buffer and boiled for 5 min. The samples were subjected to SDS-PAGE and western blotting with the indicated antibodies as described previously (Yoshida et al., 2011, Yoshida et al., 2015b). Two independent experiments were performed to confirm the results of pilot studies.

## Acknowledgement

The authors thank Drs. David Friedman and Jonathan Backer for suggestions. This work was supported by NIH grant GM-110215 to J.A.S.

---

**Supplementary Figure 1.**
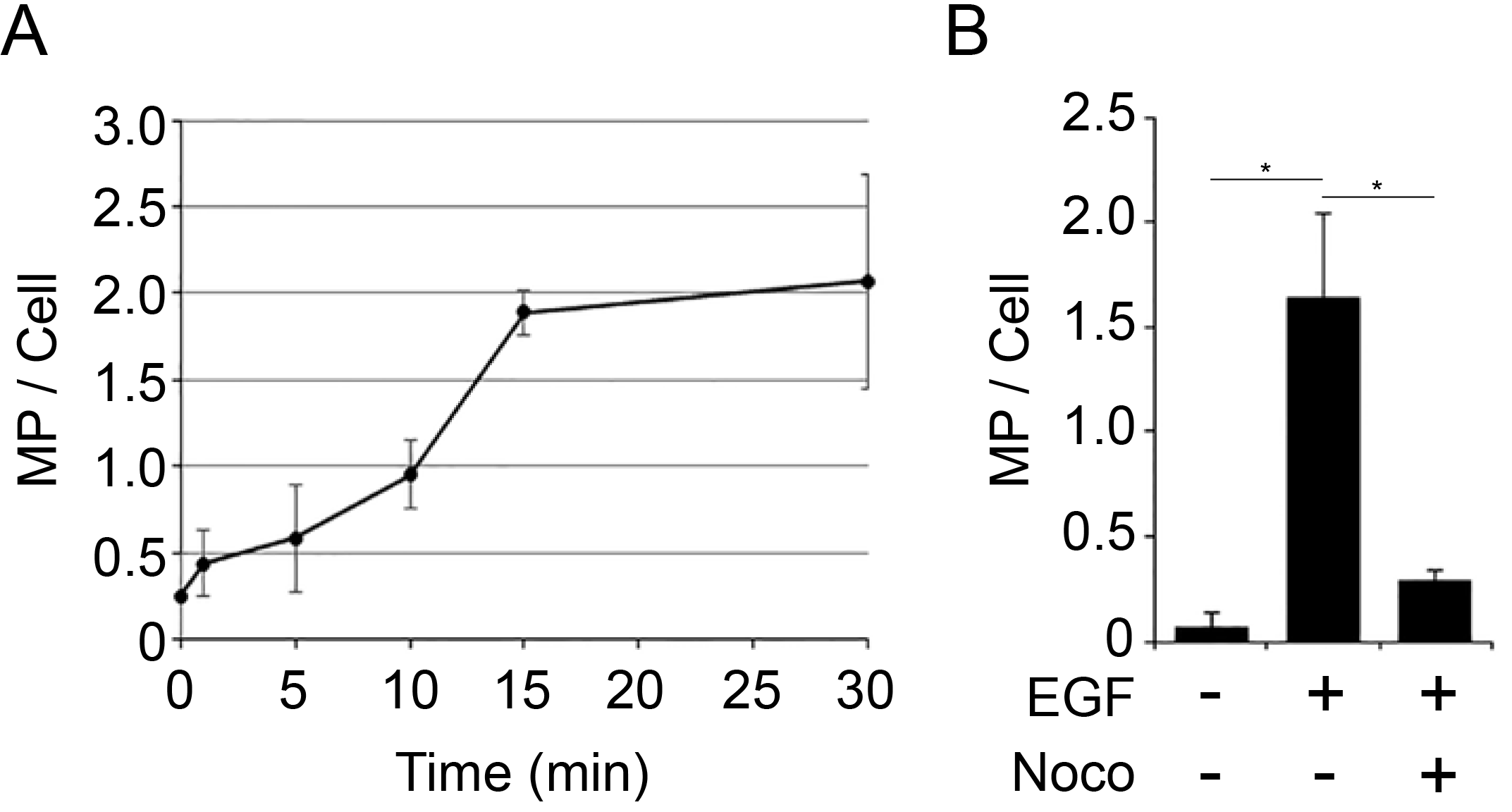
EGF-induced macropinocytosis is inhibited by nocodazole. (**A**). Time-course of EGF-induced macropinocytosis. MEF were stimulated with EGF in the presence of FDx, then washed, fixed and scored for macropinosomes. The number of macropinosome in EGF-treated MEF increased 15 min after the treatment. (**B**). Nocodazole treatment blocked EGF-induced macropinocytosis. Cells were incubated 15 min with FDx, then washed, fixed and scored for macropinosomes. More than 15 cells were observed for each condition. A one-way ANOVA was applied for statistical analysis. *p<0.05.

**Supplementary Figure 2.**
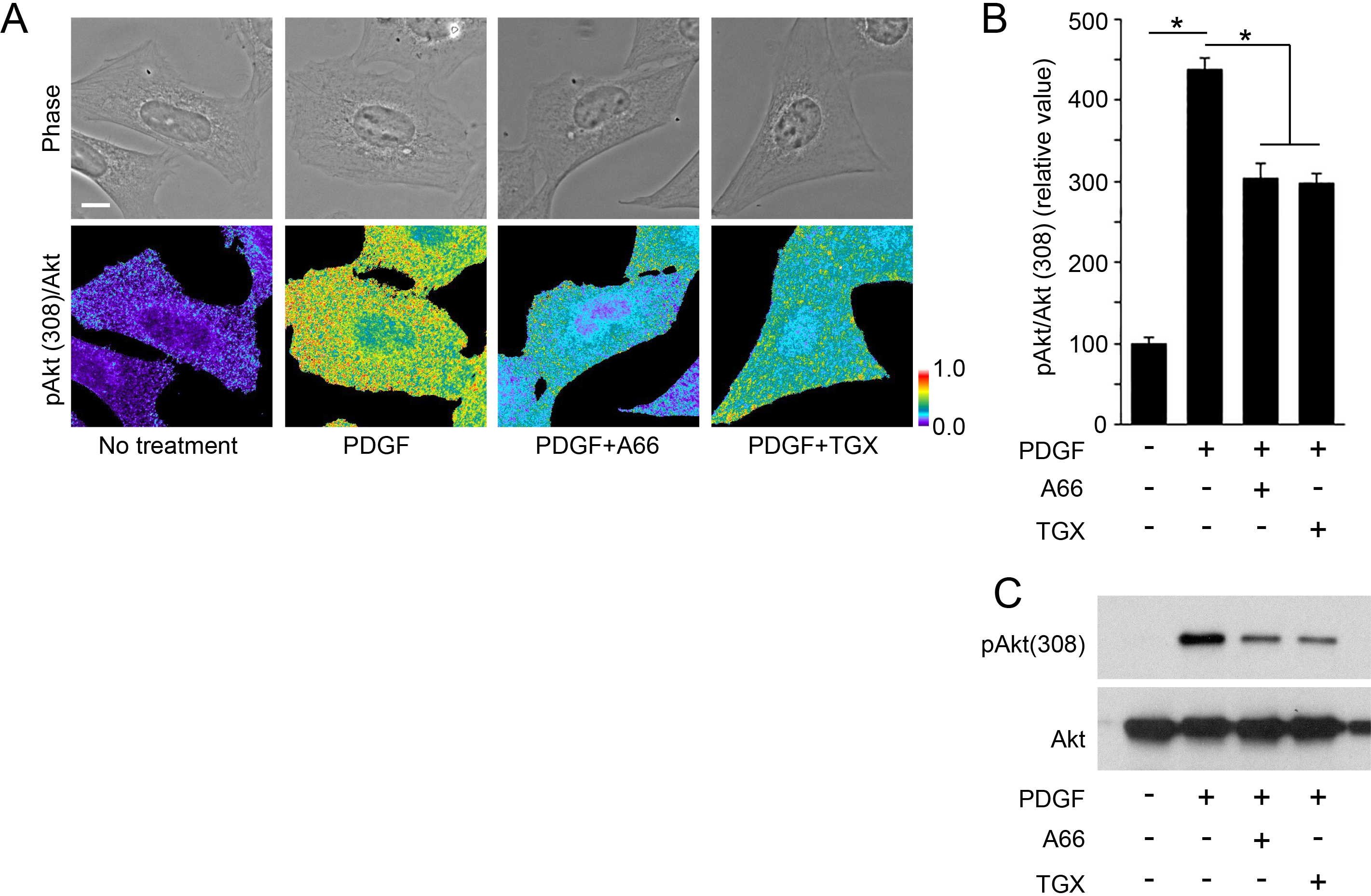
Validation of pAkt/Akt ratio imaging to quantify Akt phosphorylation in individual cells. (**A**) MEF were treated for 15 min with 2 nM PDGF with or without PI3K inhibitors A66 (p110α-specific) and TGX221 (p110β-specific). After fixation, samples were stained with anti-pAkt(308) and anti-Akt antibodies. Alexa-488-anti-rabbit and Alexa-594-anti-mouse antibodies were applied to detect pAkt(308) and Akt, respectively. Phase, pAkt(308), and Akt images were taken. Ratio images were calculated by dividing the pAkt (308) fluorescence by Akt fluorescence. Scale bar is 10 μm. Color bar indicates relative values of ratio intensities. (**B**) Quantification of ratio imaging in (A). The ratio value of the PDGF-treated sample was higher than that of no treatment, and was attenuated by A66 or TGX221. More than 10 cells were observed for each condition. A one-way ANOVA was applied for the statistics. *p<0.05. (**C**) For biochemistry, cell lysates were prepared in the same conditions as in (A). Western blot analysis showed that PDGF treatment induced Akt phosphorylation, which was attenuated by A66 and TGX221.

**Supplementary Figure 3.**
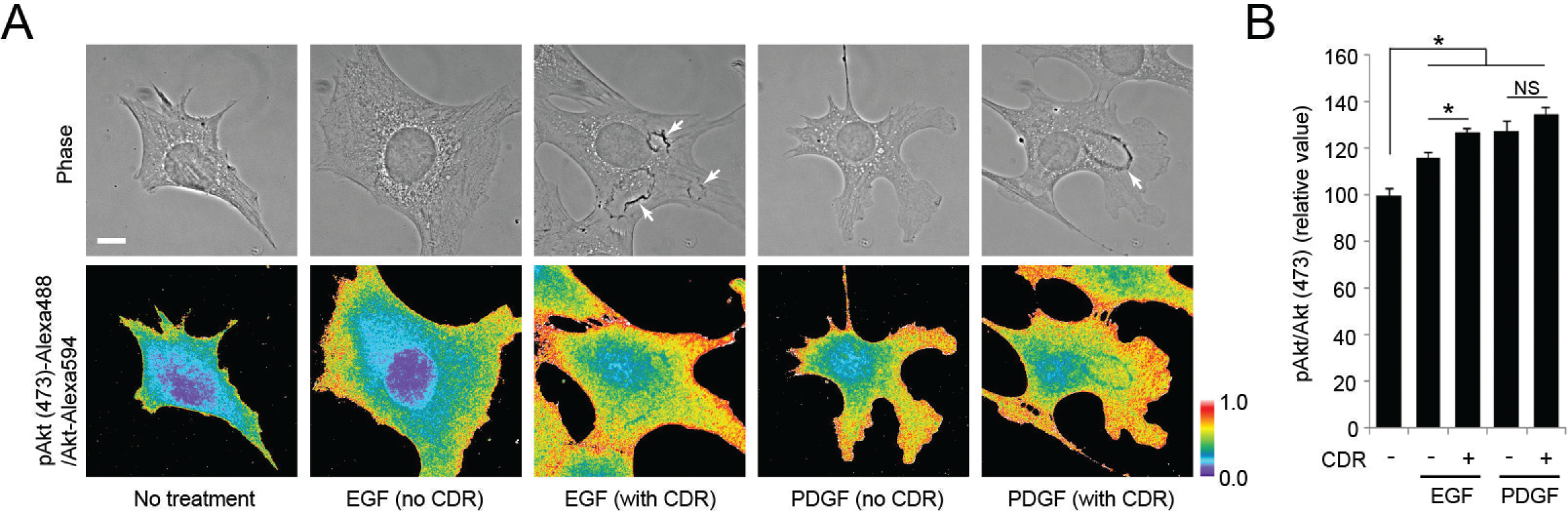
(**A**) Phase and ratio (pAkt(473)/Akt) images for MEF in different conditions, as in Figure 4. Scale bar is 10 μm. Color bar indicates relative values of ratio intensities. (**B**) Quantification of ratio imaging in (A). The relative ratio values of EGF-stimulated cells showing CDR (EGF, CDR+) were significantly higher than those of EGF-stimulated cells showing no CDR (EGF, CDR-). A one-way ANOVA was applied for the statistics. *p<0.05. More than 10 cells were observed for each assay. *p<0.05.

